# A neuronal signature for monogamous reunion

**DOI:** 10.1101/675959

**Authors:** Jennifer L. Scribner, Eric Vance, David S.W. Protter, William M. Sheeran, Elliott Saslow, Ryan Cameron, Eric Klein, Jessica C. Jimenez, Mazen A. Kheirbek, Zoe R. Donaldson

## Abstract

Pair bond formation depends vitally on neuromodulatory signaling within the nucleus accumbens, but the neuronal dynamics underlying this behavior remain unclear. Using in vivo Ca^2+^ imaging in monogamous prairie voles, we found that pair bonding does not elicit differences in overall nucleus accumbens Ca^2+^ activity. Instead, we identified distinct neuronal ensembles in this region recruited during approach to either a partner or novel vole. The partner-approach neuronal ensemble increased in size following bond formation and differences in the size of approach ensembles for partner and novel voles predicts bond strength. In contrast, neurons comprising departure ensembles do not change over time and are not correlated with bond strength indicating that ensemble plasticity is specific to partner approach. Further, the neurons comprising partner and novel approach ensembles are non-overlapping while departure ensembles are more overlapping than chance, which may reflect another key feature of approach ensembles. We posit that the features of the partner approach ensemble and its expansion upon bond formation make it a potential key substrate underlying bond formation and maturation.

**Highlights:**

- We performed in vivo Ca^2+^ in the nucleus accumbens of pair bonded prairie voles
- Overall nucleus accumbens activity did not differ during partner versus stranger interaction
- Distinct approach neurons exist for the partner and for the stranger
- Partner-approach ensemble increases as partner preference emerges
- We identify a putative neuronal substrate underlying bond formation and maturation

---

In fewer than 10% of mammalian species, humans included, individuals form mating-based pair bonds (Kleiman, 1977; Lukas and Clutton-Brock, 2013). Pair bonds are maintained and reinforced over time by a selective desire to seek out and interact with a bonded partner. This behavior is not exhibited by most laboratory rodents, including mice and rats. However, in monogamous prairie voles, pair bonding is easily assessed using a test in which the focal animal chooses between spending time with a pair-bonded partner or a novel opposite-sex vole tethered on opposite sides of a testing chamber (Fig 1E) (Getz et al., 1981; Williams et al., 1992). This partner preference test provides an opportunity to examine neural activity while an animal is displaying a pair bond and to assess how behavior and neural activity change as a bond matures.

**Figure 1.**
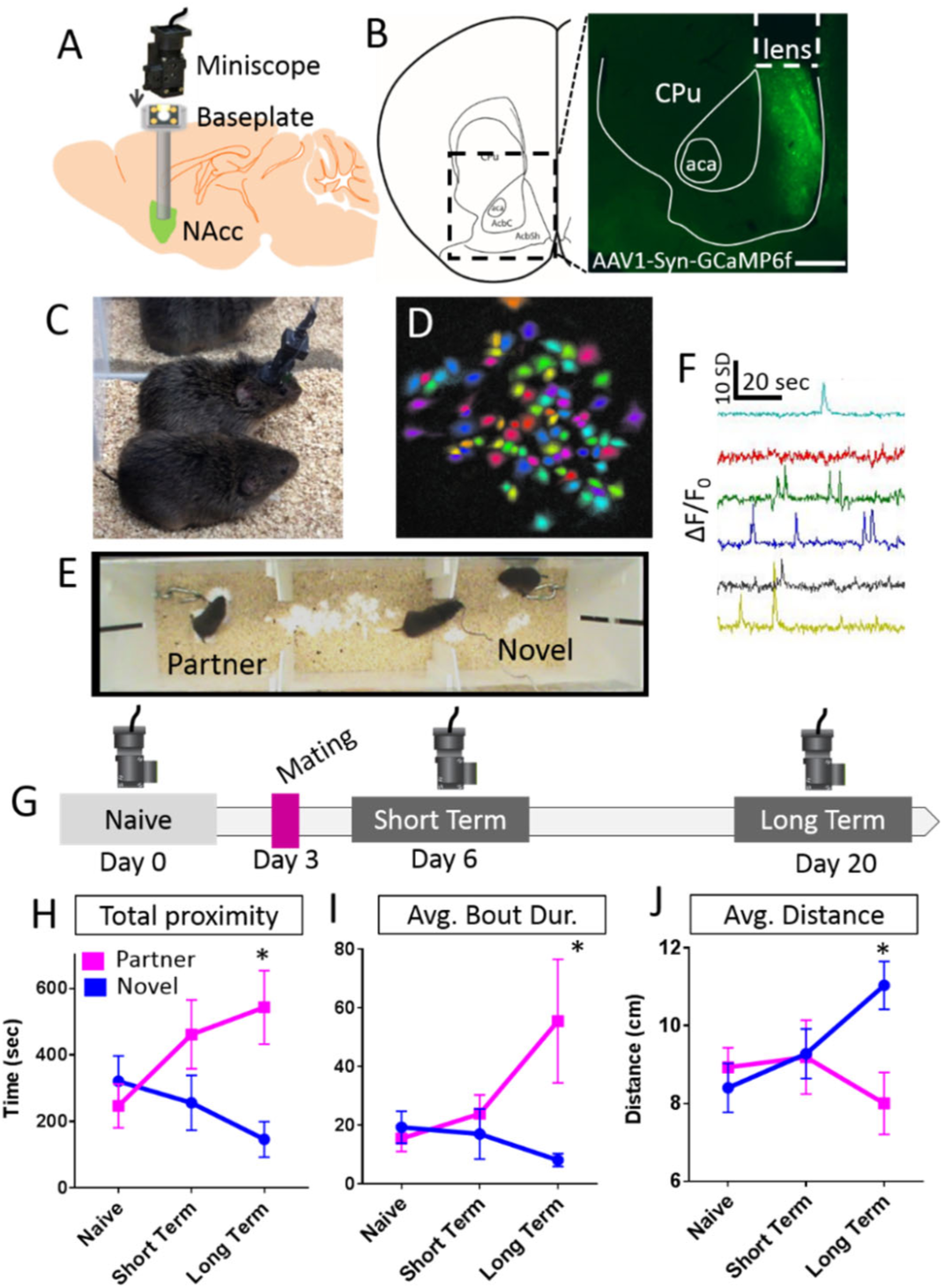
Ca^2+^ imaging in monogamous prairie voles. A) Voles were injected with AAV1-hSyn-GCaMP6f and a GRIN lens was implanted into the NAc. After recovery, a baseplate was permanently affixed to the skull to enable placement of the miniscope. B) Expression of GCaMP6f and lens insertion site. Scale bar = 500 μm. C) Two voles huddling during an imaging session. D) Putative neurons identified within the FOV of one animal. E) Imaging sessions were carried out during a 20 minute partner preference test. The test animal (center) with scope attached could freely move between two opposite sex animals tethered at either end of the apparatus. F) dF/F traces for six putative neurons. G) Experimental timecourse showing imaging sessions undertaken in sexually naïve individuals (day 0) and at short (day 6) and long term (day 20) time points after partner introduction and mating. Animals cohabited with their partner continuously beginning on day 3. H–J) Differences in social choice behavior emerged during the long-term imaging session. Test animals spent more time in proximity (<10 cm) (H), exhibited longer interaction bouts (I), and were physically closer to their mating partner (J) during the long-term imaging time point but did not exhibit differences in these metrics at the naïve or short-term imaging sessions. All error bars are SE. Lens placements in Fig S1; additional behavioral analysis in Fig S2.

The nucleus accumbens (NAc) plays a large role in reward and motivation (Ikemoto and Panksepp, 1999), making it a likely brain region for encoding highly rewarding pair bonds (Walum and Young, 2018). When participants in an fMRI study thought that they were holding hands with their pair-bonded partner, rather than an unfamiliar individual, they exhibited enhanced blood oxygenation (BOLD) signal in the NAc (Kreuder et al., 2017). In prairie voles, disruption of neuromodulatory signaling within this region impairs bond formation (Walum and Young, 2018), and subsequent gene expression changes contribute to bond maintenance (Aragona et al., 2006; Resendez et al., 2016a). However, despite substantial evidence that the NAc plays a primary role in encoding pair bonds, the neuronal dynamics underlying this process and how they change as a bond progresses remain unexplored. Thus we performed in vivo Ca^2+^ imaging in the NAc of freely behaving prairie voles before and after they mated to gain insight into how pair bond formation and maturation are represented in the brain.

## Results

### Optimization of in vivo Ca2+ imaging in prairie voles

*In vivo* Ca^2+^ imaging is tractable in prairie voles. We used microendoscopes, in combination with virally-delivered synapsin-driven GCaMP6f (Resendez et al., 2016b), a fluorescent Ca^2+^ indicator (Fig 1A– E, lens placements in Fig. S1) to image NAc neuronal activity in freely moving prairie voles. Our final dataset consisted of 17 voles (7 M, 10 F). We implanted lenses in 30 voles, 26 of which has observable fluorescence four weeks post-surgery. Of these, animals were excluded for the following reasons: lens placement outside of NAc (n = 1), < 5 detectable cells (n = 1), lack of detectable partner preference during either partner preference test (n = 3; see below), occluded field of view/motion artifact/technical problems (n = 4).

On average, we identified 43 +/-20 cells per animal per imaging session (range = 5 – 117). A repeated measures ANOVA showed that the average number of cells did not change across three imaging sessions spanning 20 days (F_(2, 38)_ = 0.034, p = 0.967). We took the average ΔF/F_0_ for each cell for the first 60 seconds and the last 60 seconds of the imaging sessions. During these epochs, the animal was housed alone with no access to stimuli, serving as a measure of baseline activity. There was no significant difference in the average ΔF/F_0_ between the baselines at any of the imaging timepoints (naïve: F_(1,614)_ = 2.506, p = 0.114; short-term: F_(1,595)_ = 0.125, p = 0.724; long-term: F_(1,639)_ = 3.246, p = 0.072) We combined these two epochs to generate a single baseline measure of ΔF/F_0_ per animal per timepoint.

### Partner preference is evident during imaging sessions

Prairie vole pair bonds are hallmarked by a selective preference to interact with a monogamous partner in a partner preference test. We imaged prairie voles in a 20-minute partner preference test at three time points spanning pair bond formation: when test animals were sexually naïve (Day 0), at a short-term time point following mating and cohabitation (Day 6), and again at a long-term time point following mating and cohabitation (Day 20) (Fig 1G). As expected, sexually naïve animals did not exhibit a preference prior to mating, and we observed the emergence and strengthening of partner preference following mating and cohabitation. Specifically, we found that test voles spent more time near their partner than the stranger (percent partner interaction) by the long-term time point (Fig 1, Fig S2D; one-way t-test relative to null value of 50%: naïve: t_16_ = −0.902, p = 0.381; short-term: t_16_ = 0.928, p = 0.367; long-term: t_16_ = 2.958, p = 0.009). Accordingly, the amount of time spent with the partner or novel animal was highly correlated with chamber time (Table S1). Test animals also exhibited longer periods of interaction (interaction bouts) with their partner than with the stranger following mating (Fig 1I, Fig 2, Fig S2C; repeated measures ANOVA: Timepoint: F_(2, 26)_ = 0.419, p = 0.662, Tethered vole: F _(1, 13)_ = 6.358, p = 0.026, Timepoint X Tethered vole: F_(2, 26)_ = 2.633, p = 0.091; paired t-test – naïve: t_16_ = 0.617, p = 0.546; short term: t_14_ = −2.247, p = 0.041; long term: t_15_ = −2.766, p = 0.014). Finally, we also examined the average distance between the test animal and tethered animal when the test animal was in the partner or novel chamber. We found that after mating and cohabitation, the test animal was closer to its partner than to the novel animals when it was in the respective chamber (Fig 1J; repeated measures ANOVA: Timepoint: F_(2, 26)_ = 0.172, p = 0.843, Chamber: F _(1, 13)_ = 5.163, p = 0.039, Timepoint X Chamber: F_(2, 26)_ = 2.959, p = 0.068; paired t-test – naïve: t_16_ = 0.617, p = 0.414; short term: t_14_ = −0.805, p = 0.434; long term: t_16_ = −3.156, p = 0.006). Thus, overall interaction time, average social bout duration, and distance from stimulus animal while in the chamber all reflect the formation of a partner preference. These data also indicate that longer cohabitation leads to stronger bonds, enabling us to ask whether bond strength is represented in patterns of NAc neuron activity.

**Figure 2.**
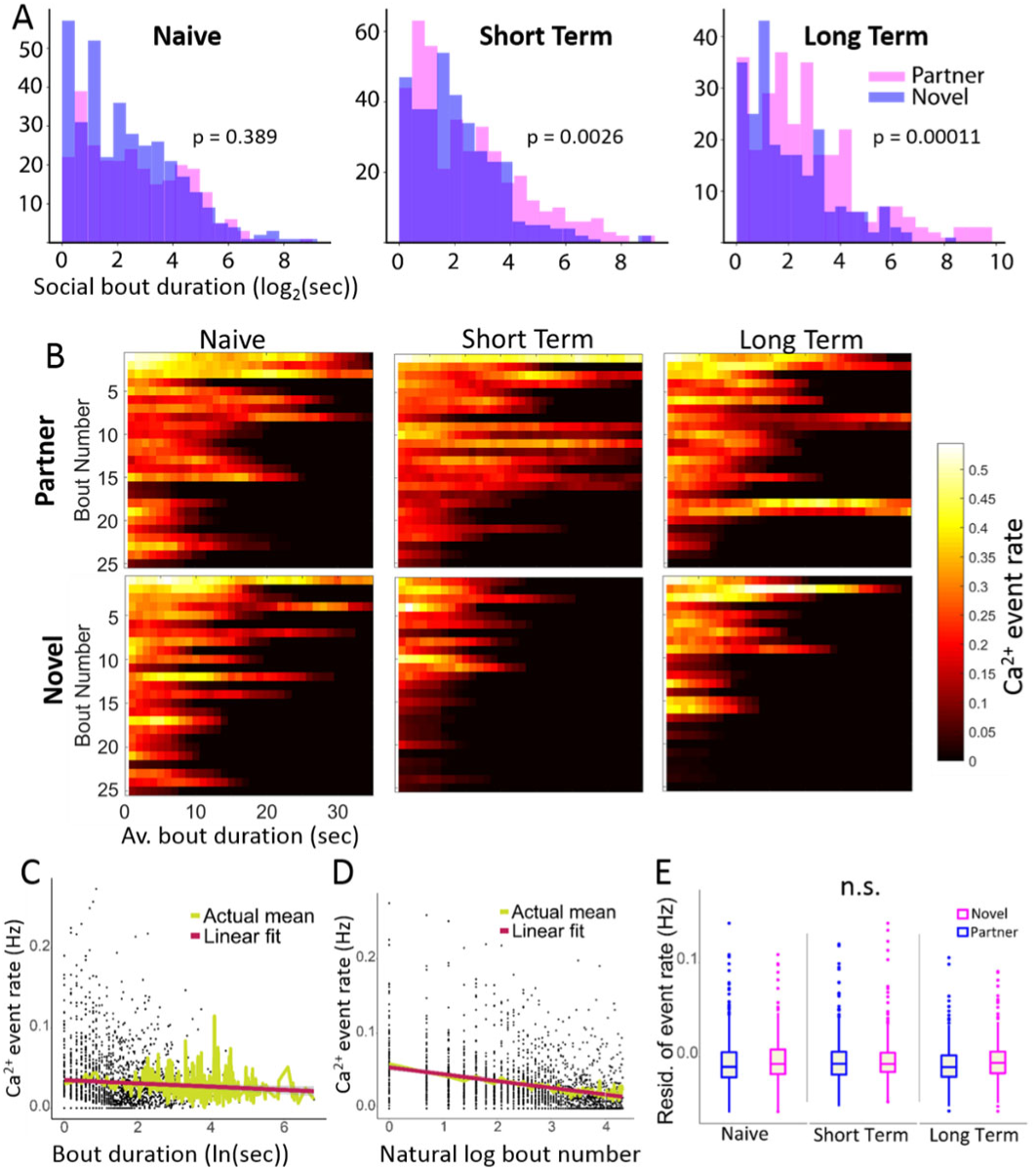
Intensity of Ca^2+^ activity across interaction bouts with different tethered animals pre- and post-bonding. A) Histograms showing frequency of social interaction bout lengths with partner (pink) and with novel (blue) animals. After mating/cohabitation, test animals had longer interaction bouts with their partner than with the novel individual. B) Heat plots show the average Ca^2+^ transient events (Hz) pooled across all cells and animals. Rows for social interaction bouts represent the average across all animals and were truncated at 35 seconds and 25 bouts, or when data was available for fewer than three animals. Bout length decreased across the test session and did not differ between stimulus animals during the sexually naïve imaging session. In contrast, test animals exhibited longer and more interaction bouts with their mating partner than the novel individual during the short- and long-term imaging sessions. C) Ca^2+^ activity was greater in short bouts than long bouts and corresponded with a logarithmic decrease in activity as bout length increased. D) Activity was greater during initial interaction bouts within a session and exhibited a logarithmic decrease across subsequent social interaction bouts. E) After controlling for differences in bout number and bout length, there were no significant differences in average calcium activity during partner and novel interaction. Residuals are plotted after regressing firingrate∼log(duration)+log(boutnumber). Individual data plotted in Fig S3.

Because we were concerned that the brevity of the imaging sessions and attachment of the scope might weaken our ability to detect partner preference, we also performed a traditional three-hour partner preference test without the scope attached following the short- and long-term imaging sessions (Days 7 and 21). We observed a partner preference at during the 3-hour test at both time points (Figure S2E; one-way t-test relative to null value of 50%: short-term: t_19_ = 2.175, p = 0.043; long term: t_19_ = 2.536, p = 0.020). The percent of time interacting with the partner was correlated across the two tests (Spearman non-parametric test, ρ = 0.537, p = 0.015), while this was not observed in the short-term and long-term imaging sessions (Spearman non-parametric test, ρ = −0.083, p = 0.751). Three test animals showed a consistent preference for the novel animal in both of the 3-hour tests; thus we excluded these animals (445, 566, and 589) from subsequent analyses. Unlike the 20-minute test, partner preference was evident even at the short-term time point in the 3-hour test (Fig S2D, E), and partner interaction was correlated across the two tests. This suggests that that the traditional, 3-hour partner preference test is more sensitive and consistent than the 20-minute version employed during imaging sessions. Overall, these data confirm the emergence and strengthening of a partner preference following mating and cohabitation.

### Population activity does not differ between partner and novel interaction

Based on human neuroimaging studies, we hypothesized that we would observe greater overall neuronal activity in the NAc while a pair-bonded partner was with its partner compared to when it was with a stranger. Unexpectedly, we found that Ca^2+^ event rate across all imaged neurons did not predict whether the test animal was interacting with its partner or the novel individual after controlling for differences in how the test animal interacted with each tethered animal. We identified all periods of social interaction during the partner preference test that were at least 1 sec in duration, referring to them as social bouts. Social bout duration did not differ in sexually naïve animals investigating two novel voles, but following mating, test animals had longer social bouts with their partner than with a novel tethered animal (Fig 2A; permutation analysis on bout duration, naïve: p = 0.389; short-term: p = 0.00255; long-term: p = 0.00011). Similar to other metrics of partner preference (Fig 1H - J), the difference in partner and novel social bout duration became more significant with longer cohabitation and mating (Fig 2A).

We found that average population activity rate was greater during initial social interaction bouts and at the beginning of a bout (linear regression, p <0.0001 for both factors; Fig 2B, C, D, Fig S3). Voles display differences in social behavior with their mate compared with a novel conspecific. Specifically, they partake in shorter social bouts during novel interactions (Fig 2A). To disentangle the effect of differences in the structure of social interactions with partner and novel tethered voles, we used a mixed-effects model to account for differences in bout duration, bout length, and individual vole variation nested within sex as a random effect to identify factors that are significant predictors of neuronal activity (Fig S3). We found that there was robust activation of the NAc during social interaction, but after accounting for differences in total number of social bouts and social bout duration, there were no differences in average activity during partner and novel interactions (Fig 2E; p = 0.061, χ^2^ = 10.535, df = 5).

Specifically, we tested three hypotheses. First, based on our behavioral data, we anticipated the largest differences in activity would be observed at the long-term time point when partner preference was most robust. We found that there was no statistically significant difference in activity when the test vole was interacting with partner and novel voles at the long term time point (p = 0.556, χ^2^ = 0.347, df = 1). Ignoring sex, we still found no significant difference in calcium event rates between partner and novel voles at this time point (p = 0.521, χ^2^ = 0.413, df = 1). We then compared rates across all three imaging sessions, combining partner interaction from the short-term and long-term imaging sessions (“partner”) and combining all interactions at the naïve time point with novel interaction during the short-term and long-term time point (“novel”). There was no statistically significant difference (p = 0.269, χ^2^= 1.224, df = 1) in event rates when interacting with partner or novel voles. Ignoring sex, we still found no significant difference (p = 0.269, χ^2^ = 1.224, df = 1). Finally, we asked whether type of interacting vole and imaging session matter, comparing six groups: Partner-naïve, novel-naïve, partner-short term, novel-short term, partner-long term, and novel-long term. This mixed effects model showed that there was no statistically significant difference (p = 0.062, χ^2^ = 10.498, df = 5) in rates between partner and novel interaction and imaging session. Ignoring sex, we still found no significant difference in rates between the 6 groups (p = 0.061, χ^2^ = 10.535, df = 5). While the natural log of social bout duration, the natural log of the bout number, the sex of the test vole, and a random effect for the individual vole itself were statistically significant predictors of event rates, the identity of the tethered vole was not statistically significant (Fig S3E). Thus overall population event rates cannot be reliably used to distinguish between partner and novel vole or the imaging session (naïve, short-term, or long-term). These findings indicate that encoding of pair bonds in the NAc does not occur via population-wide changes in activity.

### Approach cell ensemble expansion mirrors emergence of partner preference

We next asked whether activity in specific sub-populations of neurons might encode features of a pair bond. We used a Ca^2+^ event-triggered analysis to identify neurons whose transients corresponded with subsequent approach to or departure from the partner or novel animal, separately. For each cell, we calculated the median change in distance between the test animal and the tethered animal during a 1-second bin immediately following each Ca^2+^ transient (Fig 3A). We compared the observed distance change to a null model derived from randomly shuffling each cell’s transients in the partner chamber and in the novel chamber, respectively (Fig 3A). Cells with distance changes ≥ 95% of the null model were assigned as departure cells, and cells with distance changes ≤ 5%, as approach cells (Fig 3B). Therefore, each cell was assigned to a partner-associated category and to a novel-associated category of approach, departure, or neutral (Fig 3C, D).

**Figure 3.**
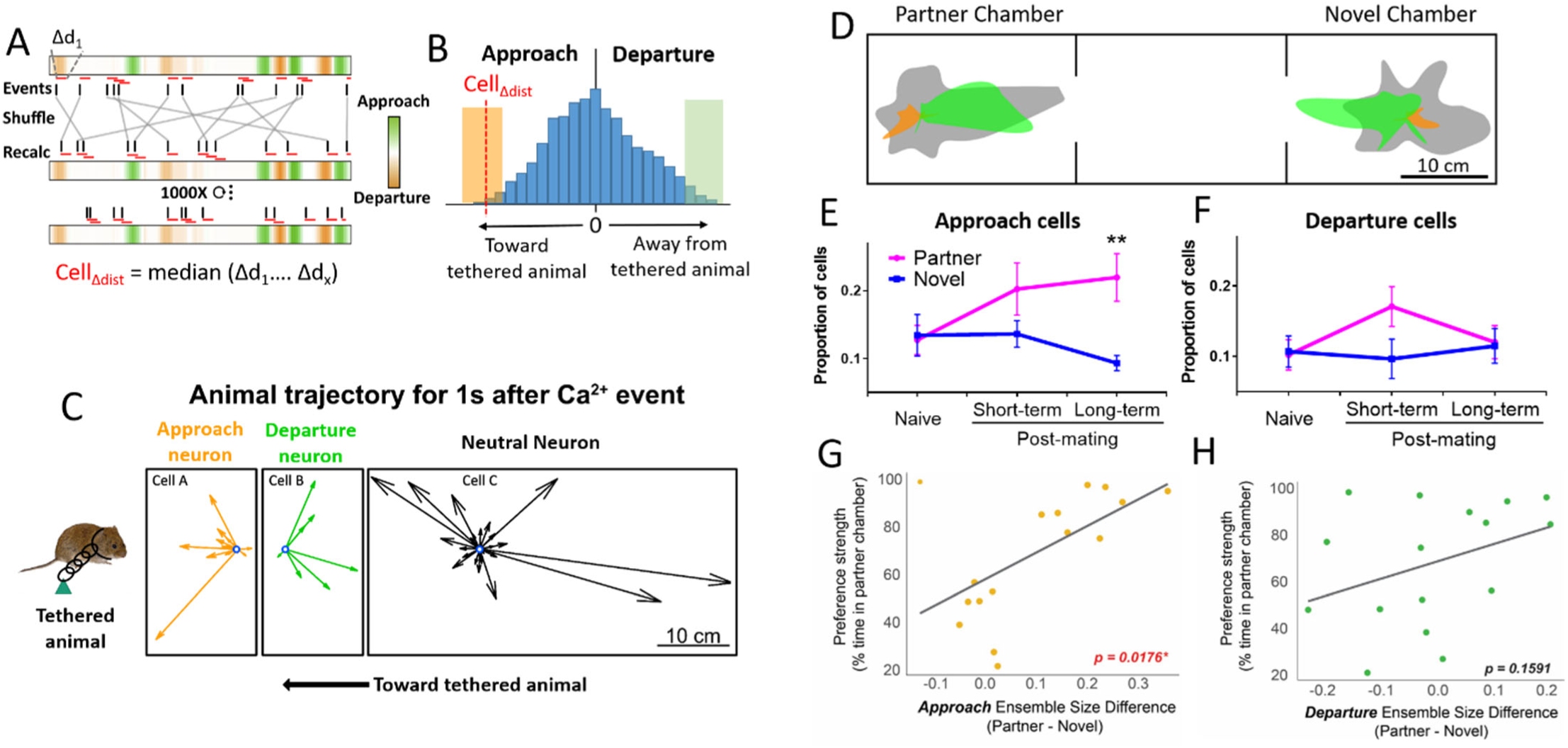
Approach cell ensemble increases upon bond maturation. A) We calculated the change in distance (Δd) between the test animal and the tethered animal in the 1 sec following each Ca^2+^ event (red rectangles) for a given cell when the animal was in the partner chamber or in the novel chamber, respectively. Events were then shuffled across distance data 1000 times to generate a null probability model. B) The median distance change was compared to a null probability distribution. Cells with an observed distance change ≥ 95% of the null distribution (green region) were defined as departure cells and those ≤ 5% as approach cells (orange region). C) Examples of approach, departure, and neutral neurons. Individual vectors represent change in distance from the stimulus animal during the 1 second following a Ca^2+^ transient and are plotted from the same origin. D) Aggregate vector maps for approach (orange), departure (green), and neutral (gray) cells with all transients normalized to the center of the partner or novel chamber. E) The proportion of partner and novel approach neurons does not differ at the naïve time point, but significantly more partner approach neurons were identified at the long-term imaging session (p = 0.021). F) The proportion of partner and novel departure cells did not differ at any time point. All error bars are SE. G) Differences in approach ensemble size (partner – novel) are correlated with partner preference strength at the long term time point when bonds have matured (ρ = 0.589, p = 0.018). H) In contrast, differences in departure ensemble size are not correlated with partner preference (ρ = 0.369, p = 0.159). Relationship between ensemble size and partner preference for each animal at other time points shown in Fig S4.

We observed an expansion in the number of partner approach cells that corresponds with the behavioral changes indicative of bond maturation (Fig 3E, G). There were no differences in partner and novel approach cells at the naïve timepoint, with differences emerging post-mating and becoming significant by the long-term time point. Animals were included only if they had n ≥ 10 cells with an event in the partner chamber and n ≥ 10 in the novel chamber. Twelve voles met criteria of having at least 10 cells with events in each chamber at each imaging timepoint (n = 416 – 504 cells). An ANOVA with repeated measures for imaging session and interaction partner revealed a significant main effect of interacting vole on the proportion of approach cells (sphericity assumed, F_1, 11_ = 7.252; p = 0.021), but no main effect of imaging session (F_2, 22_ = 2.989; p = 0.071) and no interaction (sphericity not met, Greenhouse-Geisser F_1.34, 14.7_ = 0.016; p = 0.419). We then used a post-hoc paired t-test to compare the proportion of partner and novel approach cells during each imaging session. Using the same cutoff for number of cells, one animal was excluded from naive, 2 animals from short-term, and 2 animals from the lon-term imaging session for failing to meet this cutoff. We identified significantly more partner approach than novel approach cells during the long-term imaging session (naïve: t_15_ = −0.771, p = 0.453; short-term: t_14_ = 1.387, p = 0.187; long-term: t_14_ = 3.626, p = 0.003).

We also examined whether the difference in partner and novel ensemble size for approach and departure cells correlated with partner preference strength. At the long term time point, when we consistently observed partner preference, the difference in size between the partner and novel approach ensembles were significantly positively correlated with partner preference strength (Fig S4; p-values in figure). The expansion of the partner approach cell ensemble indicates that that bond maturation may result in changes in how partner approach and novel approach are represented in the NAc (Fig 3E; main effect of partner identity, F_1, 11_ = 7.252; p = 0.021).

In contrast, we observed no differences in partner and novel *departure* cell ensemble size at any time point even though these cells were identified in the same permutation analysis (Fig 3F; main effect of Tethered vole, F_1,11_ = 1.106, p = 0.316; main effect of Imaging session F_2,22_ = 1.178, p = 0.327; Session X Tethered vole F_2,22_ = 1.91, p = 0.172). Likewise, differences in partner and novel departure ensemble size were not significantly correlated with partner preference strength (Fig S4, p-values in figure). Thus, differences in approach ensembles are unlikely due to an unanticipated variable because such a variable would equally affect identification of approach and departure cells.

As a final control, we also asked whether approach and departure cells were sensitive to the direction of travel. Specifically, partner approach and novel departure represent the same direction of travel relative to the apparatus. The same is true for novel approach and partner departure. If the direction of travel was the primary driver of activity within these cells, we would expect overlap between partner approach::novel departure and novel approach::partner departure ensembles. This was not the case. When we shuffled cell identities, we found that the overlap observed between approach::novel departure and novel approach::partner departure ensembles was not greater than chance (p > 0.05; Table S2).

### Approach/Departure cell transients primarily occur prior to approach/departure

We found that the majority of approach and departure cells exhibited transients prior to rather than during social approach or departure, indicating that these cells may directly modulate the decision to approach or leave the tethered animal. The method we used to identify approach and departure cells did not distinguish between cells whose events preceded transition to approach or departure versus those where events occurred during ongoing approach/departure. To determine whether approach-cell transients primarily occur during or prior to approach, we carried out the same permutation analysis but calculated the change in distance between test and stimulus animal for the one second ***prior*** to a Ca^2+^ event to identify cells in which transients consistently occurred after the test animal had already initiated approach. We found that on average 26.3% (range: 17.4–30.7%) of approach cells and 38.8% (range: 25– 47.4%) of departure cells represented cells whose Ca^2+^ events occurred while the animal was already approaching or departing (Table S3). For the remaining cells, approach began after the transient, suggesting a potential behavioral transition.

### Approach ensembles are distinct while departure ensembles overlap

Are partner and novel approach cells distinct populations? We performed a permutation analysis in which we shuffled cell identities to calculate the distribution of potential overlap among different functionally-defined populations. By comparing to this null distribution, we found that partner and novel approach neurons did not overlap more than would be expected by chance (Fig 3G, S5; naïve, p = 0.23; short term, p = 0.79; long term, p = 0.25), suggesting that partner and novel approach are independently represented in separate ensembles even prior to bonding. Somewhat surprisingly, partner and novel departure cells overlapped more than would be expected by chance (Fig 3H, S5; naïve, p = 0.004; short term, p = 0.014; long term, p = 0.019).

### Approach and departure ensembles lack topographical organization

The spatial organization of a cellular ensemble can provide insight into both important encoding properties of that network, as well as the inputs shaping its activity (Yuste, 2015). Both clustered and distributed cellular organization has been observed for networks involved in higher order cognitive processing. For instance, grid cells display functional micro-organization and clustering within the medial entorhinal cortex (Gu et al., 2018; Heys et al., 2014). However, in mouse brain areas projecting to the NAc, including the medial prefrontal cortex, amygdala, and hypothalamus (Walum and Young, 2018), ensembles encoding social information seem to lack meaningful spatial organization (Kingsbury et al., 2019; Li et al., 2017; Remedios et al., 2017). Thus, we asked whether approach and departure ensembles displayed spatial organization within NAc.

We calculated the distance between each possible cell pair of the same identity within each animal at the long-term time point (i.e., when ensembles are most robust) and compared these values to the distances between all cells that did not meet identity classification criteria (non-classified cells) (Fig. 4C). Novel approach, novel departure, and partner departure cell pairs were all found to have distance distributions that did not differ significantly from that of non-classified cells (Kolmogorov-Smirnov test, p > 0.05). Partner approach cell pair distances did differ significantly, but these pairs tended to be further apart rather than closer together (p < 0.001), suggesting a highly distributed organization. To further confirm that smaller-scale organizational patterns, such as spatially segregated clusters, did not exist in NAc, we also calculated the distance between the *closest* cell pair of the same identity for a given imaging field of view. We performed a similar permutation analysis as above in which we randomly shuffled the identities of every cell from a given animal at the long-term time point and recalculated the distance between nearest-neighbor cells of the same identity (Fig. 4D). A null distribution was created for each within-identity comparison by compiling the results of 1000 shuffles. Only 6.81% (3/44) nearest neighbor pairs across all animals were found to be closer than chance (≤ 5% of those in the null distribution); indeed, a higher proportion of pairs (20.54%; 9/44) was instead found to be further apart than chance (≥ 95% of those in the null distribution). Together, these results imply that novel and partner ensembles in the NAc do not display clustering and instead exist in a spatially distributed pattern.

**Figure 4.**
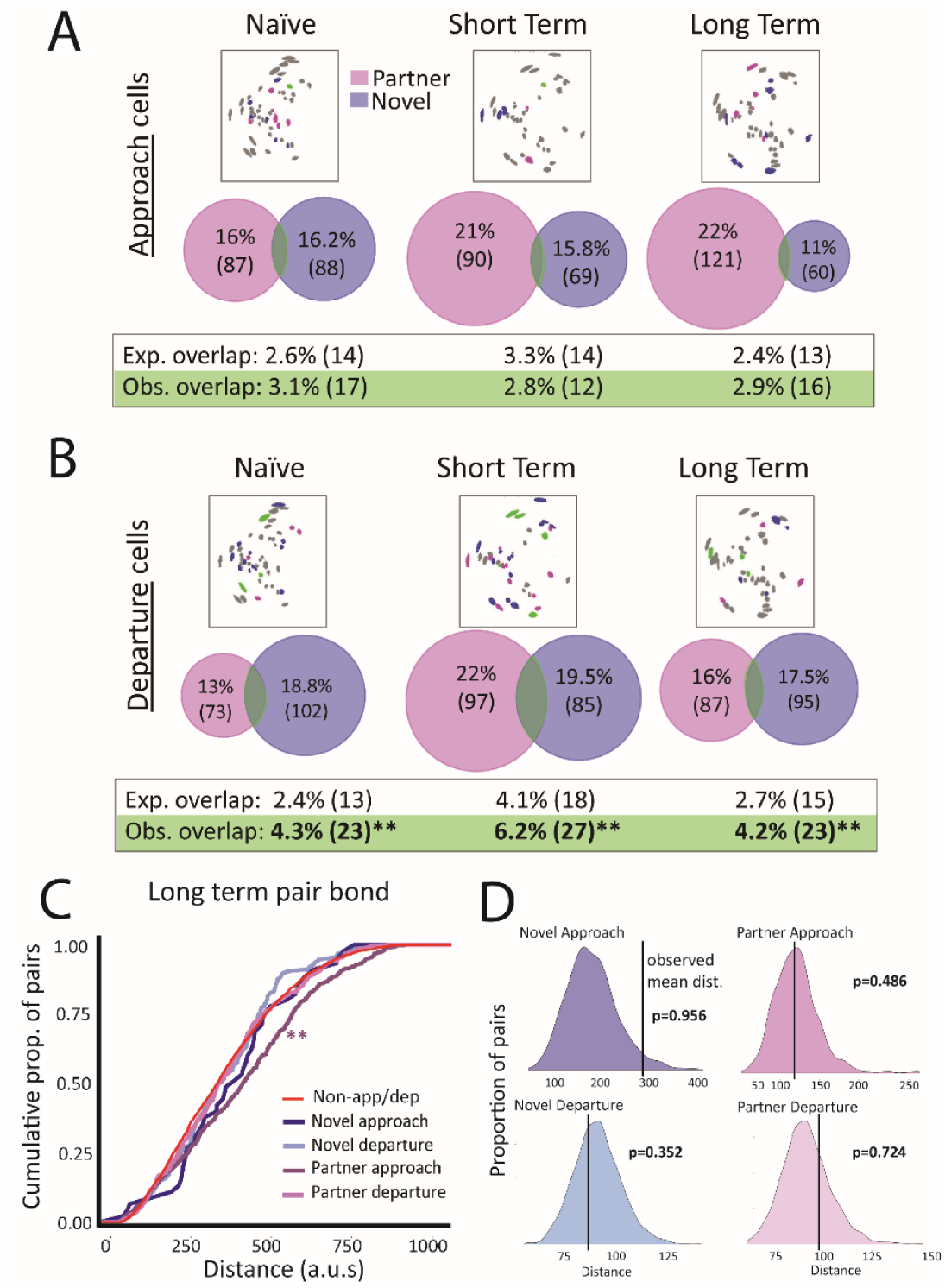
Approach and departure ensemble characteristics. A) The proportion of cells that belong to both partner and novel approach ensembles is not greater than what would be expected by chance across all time points. B) In contrast, departure ensembles overlap more than would be expected by chance at all three time points (** p < 0.02). Null distributions shown in Fig S5. C) Cells of the same identity are not closer together than other cells at the long term time point. Cumulative distribution displays the proportion of within-identity cell pairs across all animals separated by a given maximal distance (x axis), compared to control pairs of cells that did not meet any classification criteria (red line) (** p < 0.001). D) Null distributions of nearest neighbor pairs with the same cell identity. Horizontal line shows observed nearest neighbor distance. Example data are from one animal showing that cells comprising approach and departure ensembles are not clustered more than would be expected by chance.

## Discussion

Pair bonding changes the brain, and our experiments have identified a neuronal substrate contributing to these changes. Surprisingly, we found that partner-associated differences in overall NAc Ca^2+^ activity are not evident in pair bonded prairie voles. Instead, specific features of pair bonds, such as the preferential desire to approach a partner rather than a stranger, may be encoded in specific neuronal ensembles. Specifically, we identified a population of neurons, the partner approach ensemble, whose activity corresponded with subsequent partner approach. We posit that the expansion of partner approach ensembles following pair bond formation may represent a mechanism for encoding key aspects of a pair bond, such as the decision to reunite with an absent pair bonded partner.

Partner-approach neurons exhibit a number of features that make them an ideal candidate for encoding pair bond-related information. First, the expansion of the partner-approach ensemble closely parallels the emergence of partner preference, and differences in partner and novel ensemble size correlate with individual differences in preference strength (Fig 3E, G). Second, most of these cells have Ca^2+^ transients prior to rather than during ongoing approach. This would be consistent with a role for these neurons in mediating the decision to approach a particular animal. Third, the partner- and novel-approach ensembles are non-overlapping, which would be expected if these cells contain information about the specific animal being approached (Fig 4A). Notably, approach ensembles are non-overlapping even at the naïve timepoint when neither tethered animal had a significantly different familiarity/valence (both were novel opposite sex voles), suggesting that identity information alone is sufficient to recruit distinct approach ensembles. Finally, these ensembles exhibit a distributed spatial organization (Fig 4C,D), a pattern similar to that found previously in upstream social ensembles in mice. This patterning is consistent with ensemble coding rather than individual cell tuning or population rate coding as a unit of computation. Together, this suggests that plasticity in the partner approach ensemble contributes to the encoding of pair bonds.

Unlike approach ensembles, partner and novel departure ensembles overlapped more than would be expected by chance across all three time points. This difference in the properties of approach and departure ensembles further supports a functionally distinct role for these two populations. A possible interpretation of these findings is that approaching the wrong animal could have deleterious consequences (e.g., from aggression). Distinct approach ensembles may be important for identifying and deciding to approach a specific individual, while that level of specificity is not necessary when departing from a social interaction.

While our results suggest that changes in ensemble coding, especially ensemble size, may contribute to pair bonding, there are a number of limitations worth noting. Establishing Ca^2+^ imaging in voles represents a significant advance, but the specific behavioral role of approach neurons remains largely speculative without functional manipulations. Unfortunately, because these cell populations are defined by activity, rather than a particular molecular genetic component or spatial location, the methods for selectively manipulating their activity remain extremely limited and untested in voles. In addition, while we monitored the same cell population across multiple weeks as bonds formed and matured, limitations of using shared equipment made it impossible to ensure that we could repeatedly return to the same focal plane. As a result, we were unable to identify and track the same neurons across this experiment, limiting our ability to determine the potential stability of approach ensembles across time.

Prairie voles are uniquely suited to address the neuronal basis of pair bonding. They display robust behavioral changes upon bond formation, many of which are well characterized from a neuroendocrine standpoint. Developing novel approaches, such as the use of Ca^2+^ imaging will be essential for uncovering how neuroendocrine mechanisms shape the neuronal mechanisms underlying monogamy-associated behaviors. Ultimately, these technologies will enable us to address many of the questions stemming directly from the results presented in this study. For instance, what information is recruiting other cells into a partner approach ensemble (e.g. motivational valence with development of a pair bopnd)? Is activity within the partner approach ensemble required for expression of partner preference or partner-directed motivation or does this ensemble’s activity simply reflect changes in activity of upstream populations that are themselves necessary contributor to these behaviors?

In sum, we have identified a potential neuronal substrate for monogamy. The expansion of partner approach ensembles upon bond maturation and their activity patterns suggest a coding mechanism for partner preference, specifically for partner reunion. More broadly, this suggests that plasticity in NAc social ensembles may contribute to species differences in sociality and that alteration in NAc social ensembles may underlie dysfunctional social attachment. Thus, further understanding of social ensembles has the potential to reveal general mechanisms underlying natural social behavior variation and pathological social behavior.

## Author Contributions and Notes

Z.R.D., M.A.K. and J.C.J. designed research, Z.R.D., J.L.S and E.K. performed research, E.V., D.S.W.P., W.M.S., E.S., and R.C. analyzed data; and Z.R.D. wrote the paper.

The authors declare no conflict of interest.

This article contains supporting information online.

## Acknowledgments

We would like to thank the animal care staff at Columbia University and University of Colorado Boulder. Additional technical assistance was provided by Saranna Rotgard, and Ashley Cunningham. Colony founder voles were provided by Larry Young. René Hen and NYSTEM Core provided access to an Inscopix miniscope. This work was supported by awards from the Dana Foundation, the Whitehall Foundation, and NIH DP2OD026143 to ZRD.

## References

Aragona, B.J., Liu, Y., Yu, Y.J., Curtis, J.T., Detwiler, J.M., Insel, T.R., and Wang, Z. (2006). Nucleus accumbens dopamine differentially mediates the formation and maintenance of monogamous pair bonds. Nat Neurosci 9, 133–139.

Getz, L.L., Carter, C.S., and Gavish, L. (1981). The mating system of the prairie vole Microtus ochrogaster: Field and laboratory evidence for pair bonding. Behavioral Ecology and Sociobiology 8, 189–194.

Gu, Y., Lewallen, S., Kinkhabwala, A.A., Domnisoru, C., Yoon, K., Gauthier, J.L., Fiete, I.R., and Tank, D.W. (2018). A Map-like Micro-Organization of Grid Cells in the Medial Entorhinal Cortex. Cell 175, 736-750.e30.

Heys, J.G., Rangarajan, K.V., and Dombeck, D.A. (2014). The functional micro-organization of grid cells revealed by cellular-resolution imaging. Neuron 84, 1079–1090.

Ikemoto, S., and Panksepp, J. (1999). The role of nucleus accumbens dopamine in motivated behavior: a unifying interpretation with special reference to reward-seeking. Brain Research. Brain Research Reviews 31, 6–41.

Kingsbury, L., Huang, S., Wang, J., Gu, K., Golshani, P., Wu, Y.E., and Hong, W. (2019). Correlated Neural Activity and Encoding of Behavior across Brains of Socially Interacting Animals. Cell 178, 429-446.e16.

Kleiman, D. (1977). Monogamy in mammals. Quarterly Review of Biology 52, 39–69.

Kreuder, A.-K., Scheele, D., Wassermann, L., Wollseifer, M., Stoffel-Wagner, B., Lee, M.R., Hennig, J., Maier, W., and Hurlemann, R. (2017). How the brain codes intimacy: The neurobiological substrates of romantic touch. Hum Brain Mapp 38, 4525–4534.

Li, Y., Mathis, A., Grewe, B.F., Osterhout, J.A., Ahanonu, B., Schnitzer, M.J., Murthy, V.N., and Dulac, C. (2017). Neuronal Representation of Social Information in the Medial Amygdala of Awake Behaving Mice. Cell 171, 1176–1190.e17.

Lukas, D., and Clutton-Brock, T.H. (2013). The Evolution of Social Monogamy in Mammals. Science 341, 526–530.

Remedios, R., Kennedy, A., Zelikowsky, M., Grewe, B.F., Schnitzer, M.J., and Anderson, D.J. (2017). Social behaviour shapes hypothalamic neural ensemble representations of conspecific sex. Nature 550, 388–392.

Resendez, S.L., Keyes, P.C., Day, J.J., Hambro, C., Austin, C.J., Maina, F.K., Eidson, L.N., Porter-Stransky, K.A., Nevárez, N., McLean, J.W., et al. (2016a). Dopamine and opioid systems interact within the nucleus accumbens to maintain monogamous pair bonds. Elife 5.

Resendez, S.L., Jennings, J.H., Ung, R.L., Namboodiri, V.M.K., Zhou, Z.C., Otis, J.M., Nomura, H., McHenry, J.A., Kosyk, O., and Stuber, G.D. (2016b). Visualization of cortical, subcortical and deep brain neural circuit dynamics during naturalistic mammalian behavior with head-mounted microscopes and chronically implanted lenses. Nat. Protocols 11, 566–597.

Walum, H., and Young, L.J. (2018). The neural mechanisms and circuitry of the pair bond. Nature Reviews Neuroscience 19, 643.

Williams, J.R., Carter, C.S., and Insel, T. (1992). Partner preference development in female prairie voles is facilitated by mating or the central infusion of oxytocin. Annals of the New York Academy of Sciences 652, 487–489.

Yuste, R. (2015). From the neuron doctrine to neural networks. Nat. Rev. Neurosci. 16, 487–497.

