## Supplementary methods and figures for "A neuronal signature for monogamous reunion"

*Supplement*

STAR methods:

**Animals**

Prairie voles were originally obtained from a colony at Emory University that had been established from wild animals collected in Illinois. They were housed in Innovive disposable rat cages with ad lib water and rabbit chow supplemented with alfalfa cubes and sunflower seeds and cotton nestlets for enrichment. They were kept at 23–26°C with a 10:14 dark:light cycle to facilitate breeding. Animals were weaned at 21 days and housed in same sex groups of 2–4 animals until initiation of the experiment. All procedures were approved by the New York State Psychiatric Institute and Columbia University IACUC committees.

**Stereotactic surgery and baseplate attachment**

Experimental animals underwent lens implantation surgery between 60 and 90 days of age. Voles were anesthetized with 1–3% isoflurane at an oxygen flow rate of 1L/min in a head fixed stereotactic frame (David Kopf, Tujunga, CA). Body temperature was maintained at 37°C using a closed loop heating pad with rectal thermometer (David Kopf, Tujunga, CA). Eyes were lubricated with ophthalmic ointment (Puralube Vet Ointment). The fur was removed from the incision site using scissors, and the wound area was disinfected with 70% ethanol and betadine. Surgical procedures for AAV injection and GRIN lens implantation were similar to those reported by Resendez et al (2016). Briefly, the scalp and any connective tissue was removed above the frontal and parietal skull plates. Three 0.5 mm guide holes were drilled—one each in the parietal plates and one in the left frontal plate—and anchoring screws were rotated into place. The head was leveled in the anterior-posterior plane, and a 0.7 mm hole was drilled at +1.6 mm AP, +1 mm ML. The dura and meninges were carefully removed from the opening, and the wound site was irrigated with saline until bleeding stopped. Absorptive spears (Fine Science Tools (FST), Foster City, CA) were used to remove small amounts of blood as needed. A Nanoject syringe (Drummond Scientific, Broomall, PA) was lowered and 96 nL of AAV1-hSYn-GCaMP6f was injected unilaterally each at -4.8, -4.7, -4.5, -4.4, -4.3 mm DV. The syringe was left in place for 10 minutes following the last infusion. After removal of the syringe, bleeding was controlled with absorptive spears. A GRIN lens (0.6 mm diameter, 7.3 mm long; Inscopix Inc, Palo Alto, CA) was slowly lowered into position via sequential steps of 0.2 mm ventral motion followed by 0.1 mm dorsal motion until reaching a final placement of 4.5 mm DV (no tissue was aspirated out of the site). The lens and screws were affixed to the skull with Loctite 454 cured with dental cement liquid. The initial layer was covered with additional Loctite with black carbon powder (Sigma), and a well was formed around the lens. Vetbond tissue adhesive was used to seal the edges of the skin, and the lens was protected with liquid mold rubber (Smooth-On). Subcutaneous carprofen (5 mg/kg) and saline (up to 3mL) were administered peri-operatively and for 2 days after surgery to prevent dehydration and for analegesia. Animals were checked daily and the protective rubber mold was replaced as needed.

Four weeks after completion of surgery, voles were checked for GCaMP expression with a miniaturized microscope using procedures previously described(Resendez et al., 2016). Animals were briefly anesthetized with 1 – 3% isoflurane with 1L/min oxygen flow in a head-fixed stereotactic frame. The protective rubber mold was removed, and a magnetic baseplate was attached to a microscope and lowered over the implanted GRIN lens to assess the field of view for GCaMP+ neurons. If GCaMP+ neurons were detectable, the baseplate was cemented into place on the existing Loctite headcap. Imaging sessions were initiated 1 – 5 days after baseplate attachment.

**Imaging sessions**

Voles were briefly anesthetized (< 10 minutes) in order to attach the miniscope to the baseplate. Imaging sessions were conducted in a 3 chambered apparatus that enabled partner preference assessment (Fig 1E). Voles were restricted to the middle chamber via plexiglass chamber dividers for 30 minutes of recovery prior to initiating imaging. During the recovery period, stimulus animals were tethered in the side chambers and allowed to habituate prior to testing. Novel and partner stimulus animals were randomly alternated to left or right sides of the apparatus.

Imaging sessions were conducted at three time points: Naïve (day 0), short-term (day 6), and long term (day 20) (Fig 1G). Animals were housed in divided cages after the imaging session conducted on day 0, and the divider was removed on day 3 to allow mating. Test animals lived with the same partner for the duration of testing. Each imaging session consisted of a baseline recording (60 sec), access to sunflower seeds (180 sec), exposure to a novel object (180 seconds; conical tube, slide box, or ceramic shoe, randomly assigned across time points), a partner preference/social choice test (1200 seconds, described below), and a final baseline session (60 seconds). Baseline, food, and object sessions occurred in the middle chamber of the social choice test apparatus, and the dividers were removed for full access to the apparatus during the partner preference test.

$Ca^{2+}$ videos were recorded with nVista acquisition software (Inscopix, Palo Alto, CA) at 15 frames per second with 66.56 ms exposure. An optimal LED gain and power was selected for each animal based on GCaMP expression in the FOV, and the same LED settings were used for each vole throughout the series of imaging sessions.

Behavior was recorded simultaneously from cameras placed above and to the side of the arena. Multi-animal tracking was performed via idTracker with manual correction of swapped IDs as needed. The coaxial cable from the microscope was painted black because the white cable caused idTracker to occasionally register the test animal as two silhouettes. Location data for each animal was downsampled to 5 fps and aligned to the imaging data.

**Partner preference test**

Partner preference testing was performed in custom plexiglass chambers divided into 3 equal size chambers (76 cm long X 18 cm wide X 30 cm deep) (Fig 1E)(Ahern et al., 2009). The testing arenas were modified from previous designs to accommodate imaging. Chamber dividers were angled to avoid snagging the coaxial cable during imaging sessions, and the chambers could be isolated by insertion of plexiglass dividers. Finally, stimulus animals could be tethered to the side wall by sliding their tether down a groove in the wall for 3-hour tests or by tethering to an eye bolt at the base of the chamber via a carabiner during imaging sessions to prevent tangling the coaxial cable with the tether. Conventional partner preference tests consisting of 3 hours with no scope attached and no experimenter in the room were carried out the day after the short-term (day 7) and long-term (day 21) imaging sessions. Animals were provided with food and water during these tests but not during 20-minute imaging sessions. For both, stimulus animals were briefly anesthetized, and a zip tie was placed around their neck. The zip tie was then anchored to the wall or base of the apparatus via fishing swivels, and stimulus animals were allowed to habituate for at least 10 minutes prior to initiation of the test. For 3-hour tests, animals were filmed from above, and multi-animal tracking was performed via idTracker with manual correction of swapped IDs as needed. In both imaging-based and traditional tests, social interaction was defined as periods in which the test animal and the tethered animal were within 10 cm of each other, a metric that correlates strongly with hand-scoring of social interaction (Fig S2B).

**Brain collection**

Upon completion of all imaging sessions, voles were transcardially perfused with 4% paraformaldehyde in phosphate buffered saline. The head was removed and post-fixed for 5 days in 4% paraformaldehyde before extracting the brain. The brain was equilibrated in 30% sucrose, sectioned in 50 µm slices using a sliding freezing microtome (Leica), and mounted on slides. Lens placement was drawn onto corresponding mouse atlas sections (see supplemental figure 1).

**Image Processing**

Image processing on the whole population was performed using Mosaic software (version 1.0.5b; Inscopix, CA). Videos were downsampled with a binning factor of 4 (16x), and lateral brain movement was corrected using the registration engine Turboreg(Ghosh et al., 2011; Ziv et al., 2013) using a single reference frame and high-contrast features in the image to shift frames with motion to matching XY positions throughout the video. Uneven borders originating from XY translation in motion correction we cropped. $\Delta F/F_0$ videos were generated using a minimum z-projection image of the entire movie as the reference $F_0$. Subsequent $\Delta F/F_0$ videos were were temporally downsamples by a binning factor of 3 (down to 5 frames per second). Putative single cell and Ca2+ signals were isolated with an automated cell-segmentation algorithm that employs independent and principle component analyses on the $\Delta F/F_0$ video(Mukamel et al., 2009). Identified putative cells were then sorted via visual inspection to select for units with appropriate spatial configuration and Ca2+ dynamics consistent with signals from individual neurons. Ca2+ transient events were then defined by large amplitude peaks with fast rise times and exponential decay via a Ca2+ event detection algorithm (parameters: tau = 200 ms, Ca2+ transient event minimum size = 6 median average duration).

**Population activity rates**

Social interaction bouts were defined as times when the test animal was within 10 cm of a tethered vole for a minimum duration of 1 second (Fig S1B, C). The overall spiking rate for each vole was calculated as the total number of spiking events for all cells per second divided by the number of cells for each bout. Subsequent curve-fitting, linear regression, and mixed-effect models were performed in R using the tidyverse, car, and lme4 packages(Bates et al., 2015; Fox and Weisberg, n.d.; R Core Team, 2017; Wickham, 2017).

**Approach and departure cells**

We used an $Ca^{2+}$ event-triggered analysis to identify cells whose event activity preceded a decrease or increase in distance from a stimulus animal, referring to these as approach and departure cells, respectively. We conducted separate analyses for events occurring in the partner chamber and in the novel chamber. For each cell, we calculated the median change in distance between the test and stimulus animal following each event. We calculated a null distribution for change in distance by shuffling the events and repeatedly calculating the median change in distance 1000 times. Approach or departure cells consisted of cells where the mean change in distance fell within the 5% tails of the distribution, e.g. with a p-value > 0.95 for departure or < 0.05 for approach. The proportion of approach or departure cells was calculated using the number of cells with at least one event in the chamber of interest as a denominator to account for differences in the number of cells with activity in each chamber. To determine whether approach or departure cells consisted of the same population of cells, we carried out a permutation analysis, limited to cells that had at least one event in the partner and in the novel chamber (naïve, n = 588; short term, n = 476, long term, n = 568). Briefly, we calculated the number of partner approach cells, the number of novel approach cells, and the number of cells that met criteria for both, using a slightly relaxed threshold of p < 0.1. We calculated a null distribution by shuffling cell classification (e.g. partner approach or novel approach etc) and calculating cell overlap 1000 times.

The vector plots of approach vs. departure cells (Fig 3) were created by assigning the test animal and the partner/novel animal to specific points in space before finding their vectors. We identified $t_0$ and $t_1$, which correspond to the time of a $Ca^{2+}$ event within the cell and the time 1 second after that event, respectively. These time points were used to generate a change in position vector $\boldsymbol{t}$, for the test animal. For the partner/novel animal we identified the midpoint of its position at those two timepoints, $p_m$. Based on $p_m$, we calculated a relative position vector $\boldsymbol{r}$, from the initial position of the test animal at $t_0$ to the position $p_m$.

Next, for the partner animals we set their arbitrary position to be in the direction $\boldsymbol{d} = [-1,0]$, directly to the left, and for the novel animals at $\boldsymbol{d} = [1,0]$, directly to the right. Based on these positions, we rotated $\boldsymbol{r}$ through an angle $\alpha$ until it pointed in the unit direction of $\boldsymbol{d}$. Finally, we rotated $\boldsymbol{t}$ through that same angle $\alpha$, thereby preserving the relationship between $\boldsymbol{t}$ and $\boldsymbol{r}$ but normalizing all events to a specific direction for comparison. We obtained one vector for each neuron by taking the component-wise average of all the event vectors in a cell, therefore yielding an average relative distance change vector for each cell in each animal in each epoch.

Aggregate vectors for each cell type (approach, departure, neutral) were visualized by assigning all of the vectors in the category into bins of 20 degrees. For each bin, we added the magnitude of each vector present and plotted that magnitude vector at the halfway point of the bin. We connected the points of the vectors and smoothed out the lines using a moving mean function.

**Spatial Analysis**

Two-dimensional location coordinates were calculated for every cell present in a given imaging field of view. A matrix containing the Euclidean distances between each possible cell pair was generated for each animal for each imaging session. Cells were labeled either by their identity category (e.g., novel approach) as determined above. Subsets of the overall distance matrix were generated to examine the distances between all possible cell pairs for a given category (e.g., all possible novel approach cell pairs).

Differences in distance between cell pairs of the same identity was investigated by computing using the Empirical Cumulative Distribution Function in the ggplot2 package in R. All possible cell pairs in every imaging session in the same category were concatenated into the same function. A function was also generated for all non-categorized cell pairs, which for this analysis used p > 0.90 or p < 0.1. A Kolmogorov-Smirnov test was performed to determine significance between the categorized and non-categorized cell functions; curves that were “left-shifted” relative to the non-categorized cell curve were considered to contain cell pairs with generally shorter intercell distances, whereas curves that were “right-shifted” were considered to contain pairs that were further apart (Fig 4C).

To further determine whether the closest cell pairs of a given identity differed from what would be expected by chance, we performed a permutation analysis. Briefly, for each cell we calculated the minimum distance to a cell of the same category (e.g., the shortest distance between two partner approach cells). For each animal at the long-term time point, we calculated the mean of this value for all cells of a given category. We next established a null distribution by shuffling the identity of both cells in every possible cell pair irrespective of category (i.e., the row and column names of our overall distance matrices) for each animal 1000 times. For every shuffled iteration, we again calculated the mean shortest distance between cell pairs of the same identity. We compared the actual observed values against the histogram of shuffled values (see Fig. 4D) and classified observed values falling within the 5% tails of the distribution as significant (i.e., p-value of >0.95 for larger than chance and <0.05 for smaller than chance). We performed this analysis for each category within each animal, excluding populations with either no observed cells or only a single cell meeting the appropriate classification criteria. All spatial analyses were performed in R.

**Data availability statement**

The data that support the findings of this study are available in figshare. For review, private links have been provided at the end of each figure caption, and a public DOI will be made available upon publication. Raw imaging and behavior data are available from the corresponding author upon reasonable request.

**Code availability**

MATLAb scripts are available from the corresponding author upon reasonable request. Code will be added to github prior to publication.

Supplementary table 1:

**Correlation between interaction and chamber time**

| | **Naïve** | **Short Term** | **Long term** |
|---|---|---|---|
| **Partner** | r = 0.973; p = 6.4 x $10^{-11}$ | r = 0.882; p = 3 x $10^{-6}$ | r = 0.983; p = 1.6 x $10^{-12}$ |
| **Novel** | r = 0.929; p = 6.9 x $10^{-8}$ | r = 0.951; p = 4.5 x $10^{-9}$ | r = 0.937; p = 3.2 x $10^{-8}$ |

Supplementary table 2:

**Proportion of cells where direction of travel was predictive of activity**

| **NAÏVE** | **Partner Approach** | **Novel Departure** | **Partner Departure** | **Novel Approach** |
|---|---|---|---|---|
| **Number of approach/departure cells** | 100 | 106 | 99 | 115 |
| **Overlapping cells** | 22 | | 12 | |
| **p-value** | 0.068 | | 0.493 | |

| **SHORT TERM** | | | | |
|---|---|---|---|---|
| **Number of approach/departure cells** | 122 | 80 | 116 | 98 |
| **Overlapping cells** | 13 | | 12 | |
| **p-value** | 0.936 | | 0.891 | |

| **LONG TERM** | | | | |
|---|---|---|---|---|
| **Number of approach/departure cells** | 114 | 62 | 148 | 102 |
| **Overlapping cells** | 16 | | 12 | |
| **p-value** | 0.947 | | 0.234 | |

Supplementary table 3:

**Proportion of cells where $Ca^{2+}$ transients occur during rather than prior to approach/departure**

| **NAÏVE** | **Partner Approach** | **Novel Approach** | **Partner Departure** | **Novel Departure** |
|---|---|---|---|---|
| **Number of approach/departure cells** | 100 | 115 | 99 | 106 |
| **Number of cells with transients during ongoing approach/departure** | 24 | 20 | 37 | 42 |
| **Proportion of cells** | 0.240 | 0.174 | 0.374 | 0.396 |

| **SHORT TERM** | | | | |
|---|---|---|---|---|
| **Number of approach/departure cells** | 122 | 98 | 116 | 80 |
| **Number of cells with transients during ongoing approach/departure** | 34 | 28 | 55 | 20 |
| **Proportion of cells** | 0.279 | 0.286 | 0.474 | 0.250 |

| **LONG TERM** | | | | |
|---|---|---|---|---|
| **Number of approach/departure cells** | 114 | 102 | 148 | 62 |

| | | | | |
|---|---|---|---|---|
| **Number of cells with transients during ongoing approach/departure** | 35 | 30 | 62 | 21 |
| **Proportion of cells** | 0.307 | 0.294 | 0.419 | 0.339 |

196

**Supplementary Figure 1.**

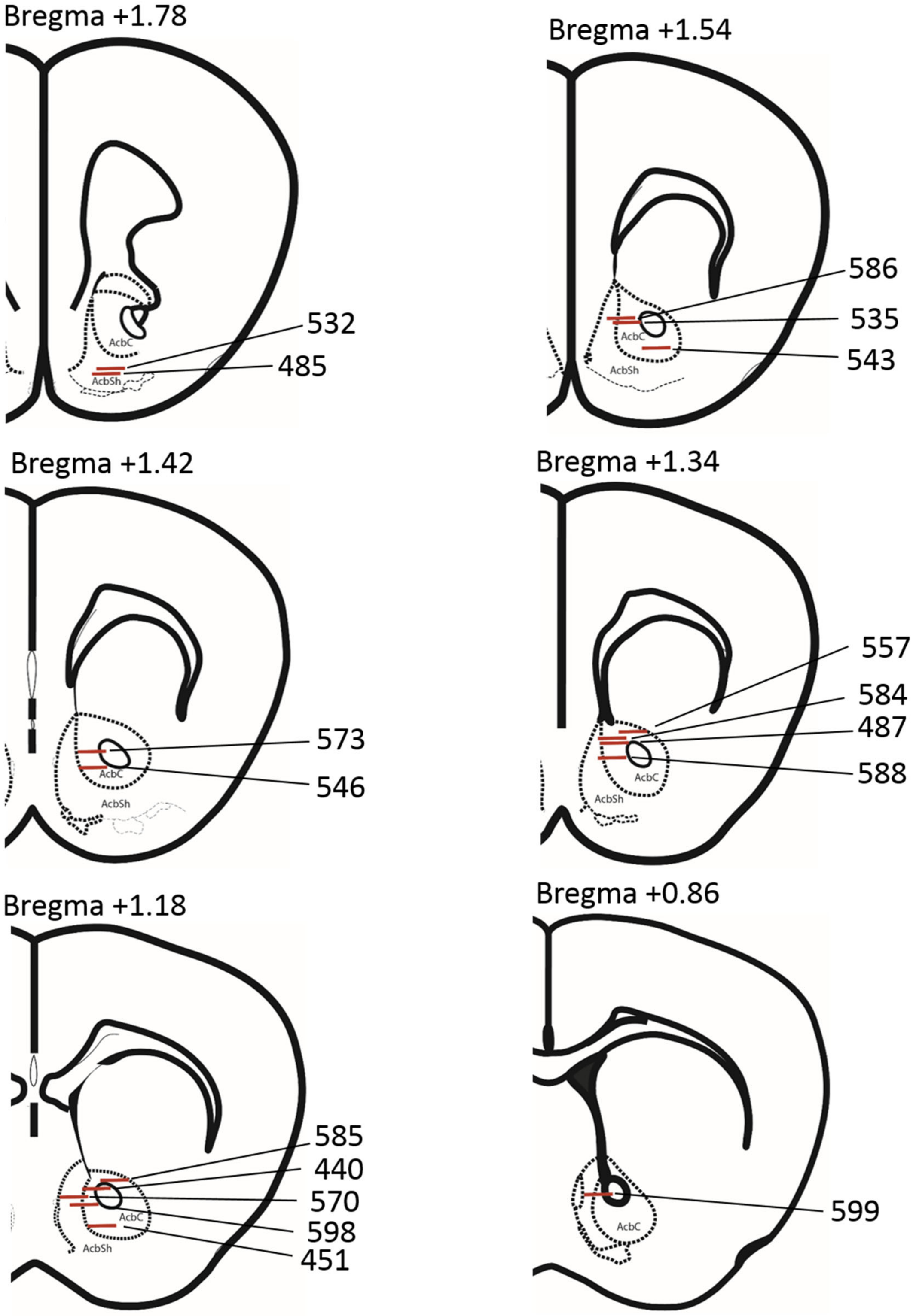

**Fig S1. Lens placements.** Adapted from mouse atlas(Franklin and Paxinos, 2008) with locations relative to bregma (mm). AcbC = Accumbens core. AcbSh = accumbens shell.

**Supplementary Figure 2.**

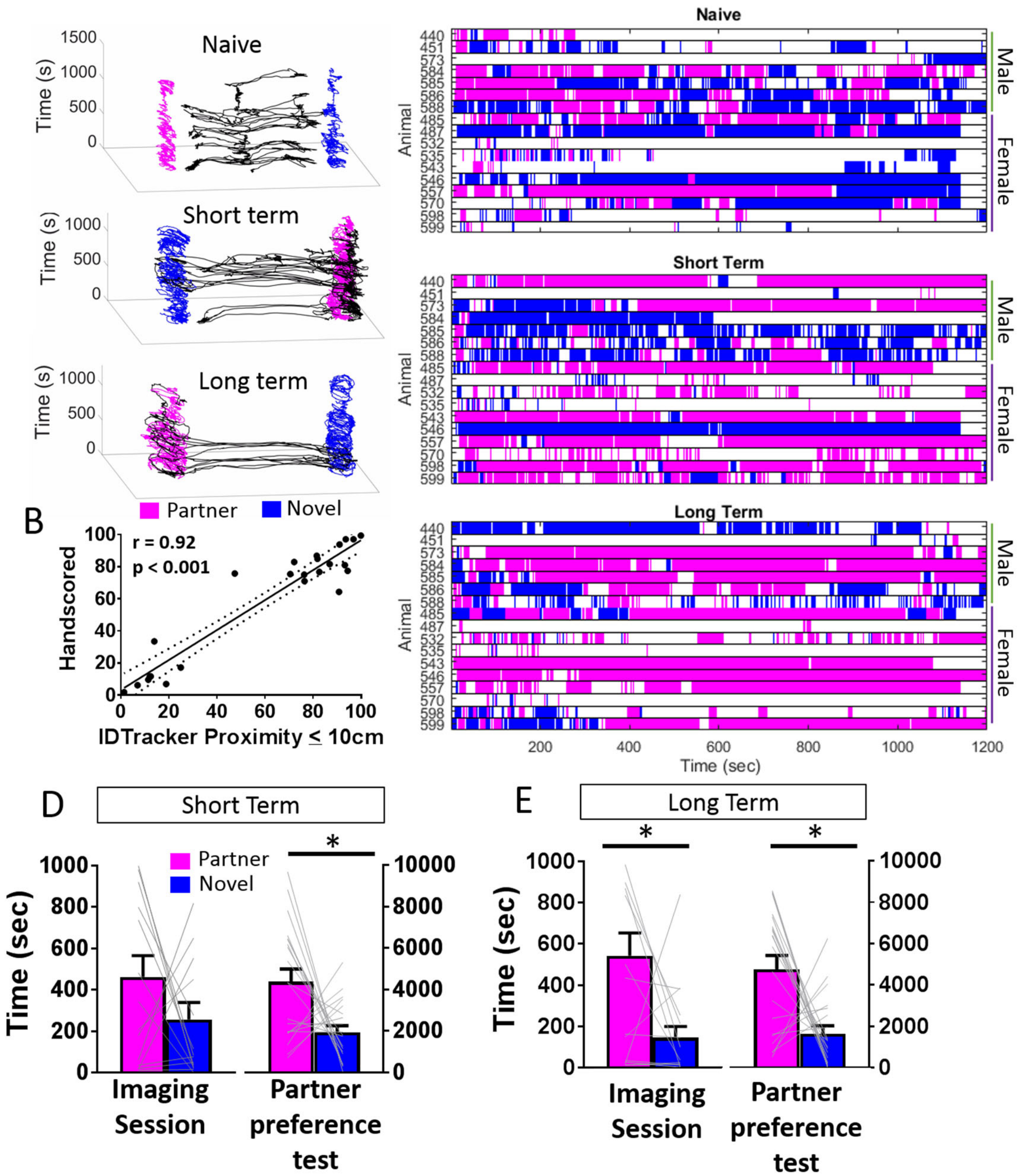


**Fig S2. Partner preference behavior.** A) Movement plots for a single test animal during each imaging session. Test animal is shown in black and can move freely between the tethered partner (pink) and the novel animal (blue). B) Comparison of hand scoring versus ID tracker. Percentage of time spent with partner is plotted. Using a cutoff of ≤ 10 cm between animals produced measures of proximity/interaction that were consistent with hand scored behavior. C) Interaction plots for each animal during each imaging session. Time with partner in pink, novel in blue. D, E) Partner preference behavior during the imaging session and a three-hour test at the short term (d) and long term (E) time points (Mean +/- SE).

**Supplementary Figure 3.**

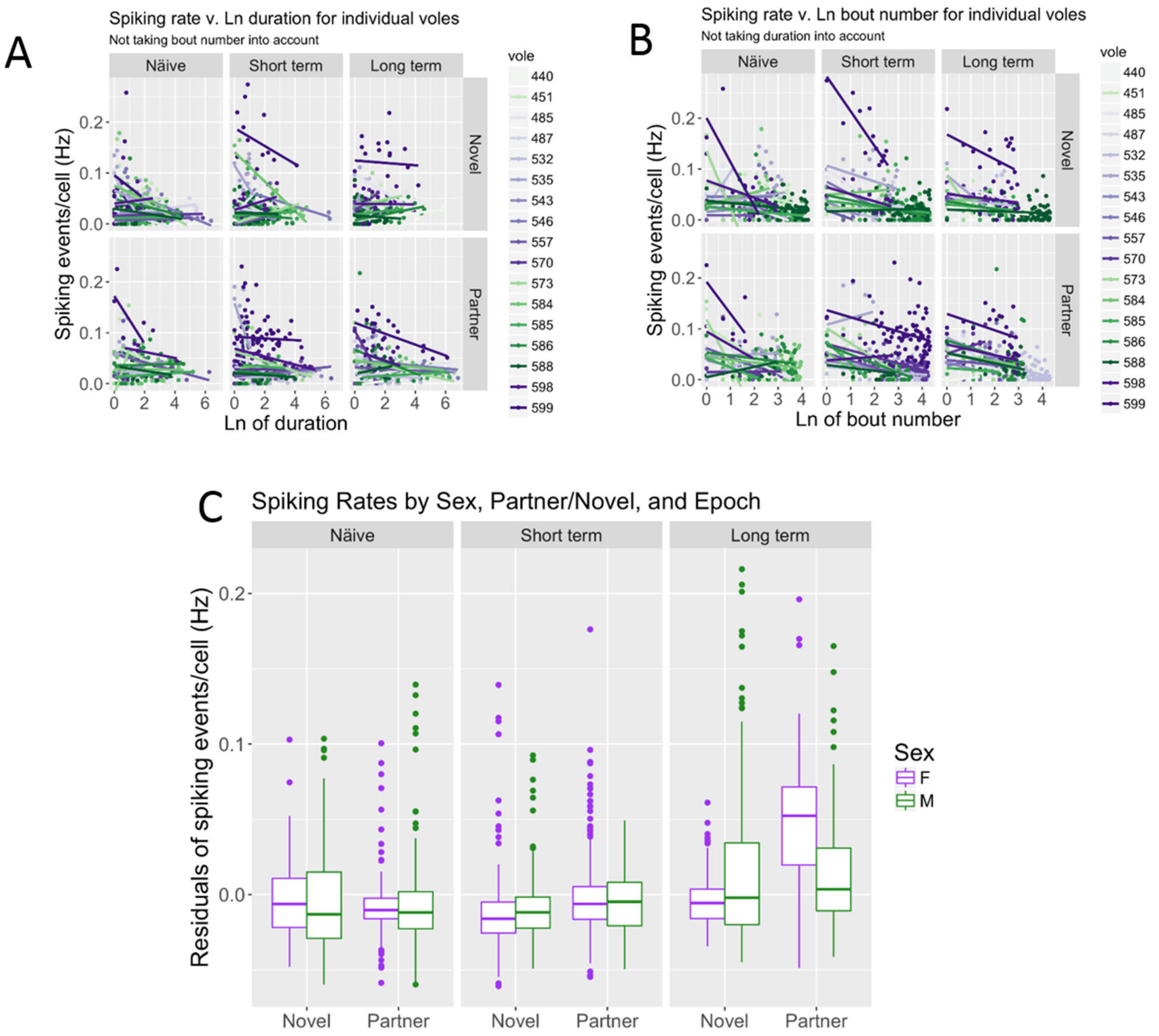


**Fig S3. Population $Ca^{2+}$ activity.** Natural log of duration and natural log of bout number were statistically significant predictors of firing rate. A and B) Relationship between Ca2+ spiking rate and ln(duration) (A) and between spiking rate and ln(bout number) (B) for individual voles. Males are plotted in green and females plotted in purple. C) The residuals of spiking rate for the linear regression model controlling for ln(duration) and log(bout number) for male and female voles interacting with partner/Novel voles in each imaging session. This difference was statistically significant ($p<0.0001$). However, much of that effect was due to female vole 599 having a much higher firing rate than the other voles. When this vole was excluded from the analysis, female voles had only a 0.002 (0.2%) higher spiking rate than male voles when interacting with tethered voles, and this difference was not statistically significant ($p=0.104$). Males and females were combined and vole 599 is excluded from figure 2E.

**Supplementary Figure 4.**

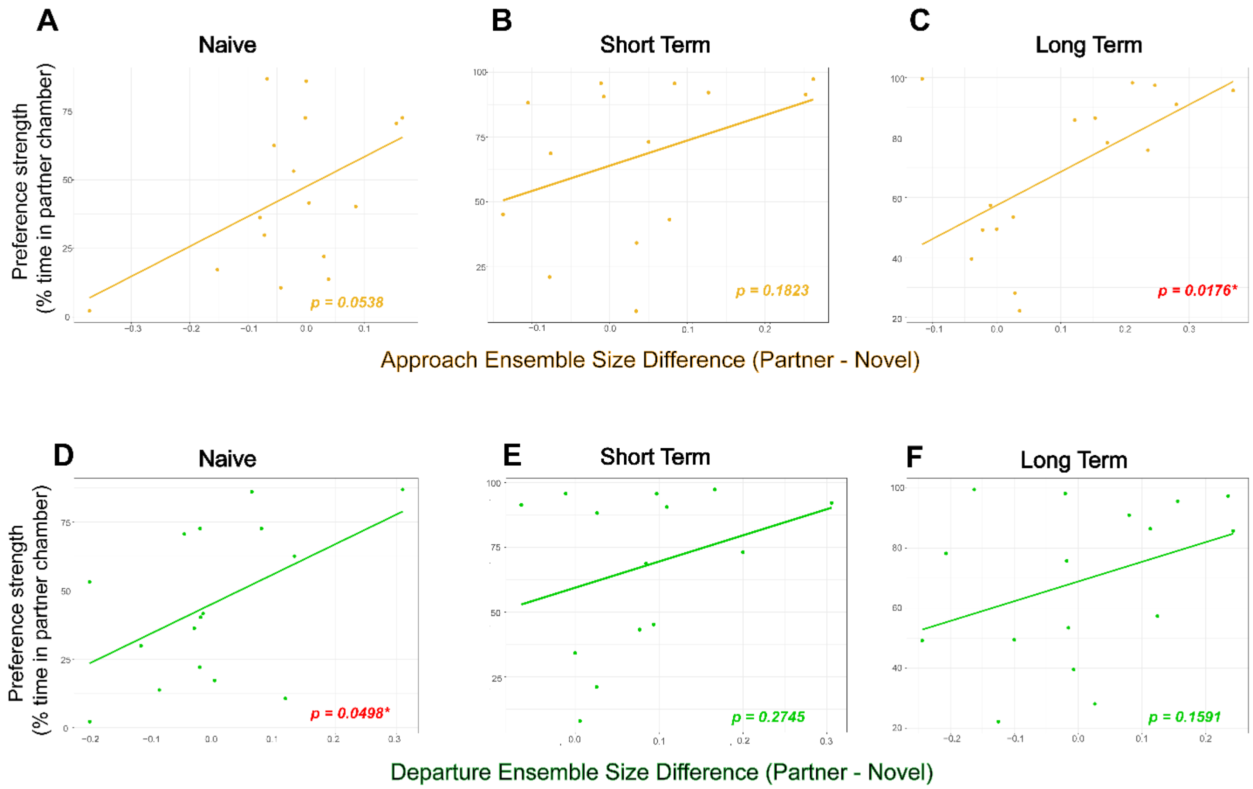


**Fig S4. Relationship between preference strength and difference in ensemble size.** We calculated the difference in partner and novel ensemble size (Approach A – C; Departure D – F) and asked whether these differences correlate with strength of partner preference. Spearman non-parametric test, Approach ensemble size difference: naive: ρ = 0.490, p = 0.054; short term: ρ = 0.378, p = 0.182; long term: ρ = 0.589, p = 0.018; Departure ensemble size difference: naive: ρ = 0.498, p = 0.0499; short term: ρ = 0.313, p = 0..275; long term: ρ = 0.369, p = 0.159).

**Supplementary Figure 5.**

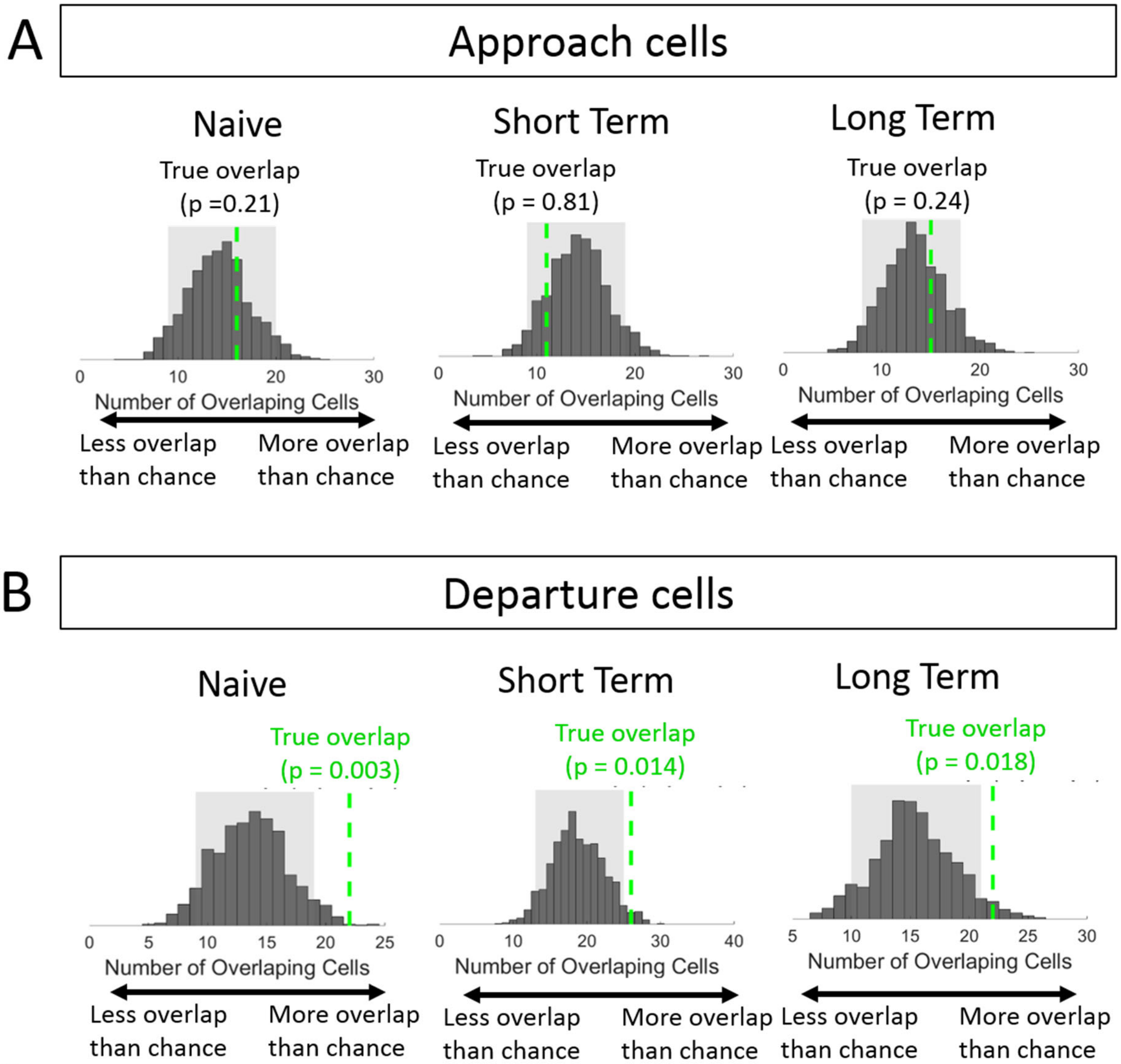


**Fig S5. Cell identity overlap between partner and novel departure cells.** Chance overlap was determined by randomly assigning cell identity and calculating overlap in 1,000 iterations to generate a probability distribution (gray). A) The true overlap (green dotted line) between partner and novel approach cells was not greater than would be expected by chance in all three imaging sessions. B) The true overlap (green dotted line) between partner and novel departure cells was greater than would be expected by chance in all three imaging sessions.

### References

Ahern TH, Modi ME, Burkett JP, Young LJ. 2009. Evaluation of two automated metrics for analyzing partner preference tests. *Journal of neuroscience methods* **182**:180–8. doi:10.1016/j.jneumeth.2009.06.010

Bates D, Mächler M, Bolker B, Walker S. 2015. Fitting Linear Mixed-Effects Models Using lme4. *Journal of Statistical Software, Articles* **67**:1–48. doi:10.18637/jss.v067.i01

Fox J, Weisberg S. n.d. An {R} Companion to Applied Regression, 2nd ed. Thousand Oaks, CA: Sage.

Franklin K, Paxinos G. 2008. The Mouse Brain in Stereotaxic Coordinates, 3rd ed. Academic Press.

Ghosh KK, Burns LD, Cocker ED, Nimmerjahn A, Ziv Y, Gamal AE, Schnitzer MJ. 2011. Miniaturized integration of a fluorescence microscope. *Nat Methods* **8**:871–878. doi:10.1038/nmeth.1694

Gunaydin LA, Grosenick L, Finkelstein JC, Kauvar IV, Fenno LE, Adhikari A, Lammel S, Mirzabekov JJ, Airan RD, Zalocusky KA, Tye KM, Anikeeva P, Malenka RC, Deisseroth K. 2014. Natural neural projection dynamics underlying social behavior. *Cell* **157**:1535–1551. doi:10.1016/j.cell.2014.05.017

Mukamel EA, Nimmerjahn A, Schnitzer MJ. 2009. Automated analysis of cellular signals from large-scale calcium imaging data. *Neuron* **63**:747–60. doi:10.1016/j.neuron.2009.08.009

R Core Team. 2017. R: A language and environment for statistical computing. Vienna, Austria: R Foundation for Statistical Computing.

Resendez SL, Jennings JH, Ung RL, Namboodiri VMK, Zhou ZC, Otis JM, Nomura H, McHenry JA, Kosyk O, Stuber GD. 2016. Visualization of cortical, subcortical and deep brain neural circuit dynamics during naturalistic mammalian behavior with head-mounted microscopes and chronically implanted lenses. *Nat Protoc* **11**:566–597. doi:10.1038/nprot.2016.021

Wickham H. 2017. tidyverse: Easily Install and Load “Tidyverse” Packages, R package version 1.1.1.

Ziv Y, Burns LD, Cocker ED, Hamel EO, Ghosh KK, Kitch LJ, El Gamal A, Schnitzer MJ. 2013. Long-term dynamics of CA1 hippocampal place codes. *Nature neuroscience* **16**:264–6. doi:10.1038/nn.3329
